# A selective S-acyltransferase inhibitor suppresses tumor growth

**DOI:** 10.1101/2024.07.18.604152

**Authors:** Jia-Ying Lee, Sebastian Dilones, Timothe Maujean, Mohammad Asad, Altaf Mohd, Noam Auslander, Donita C. Brady, George M. Burslem, Eric S. Witze

**Affiliations:** Department of Cancer Biology, Perelman School of Medicine, University of Pennsylvania; Philadelphia, 19104, USA; Department of Biochemistry and Biophysics, Perelman School of Medicine, University of Pennsylvania; Philadelphia, 19104, USA; Wistar Institute; Philadelphia, 19104, USA; Abramson Family Cancer Institute, Perelman School of Medicine, University of Pennsylvania; Philadelphia, 19104, USA

## Abstract

S-acyltransferases play integral roles in essential physiological processes including regulation of oncogenic signaling pathways. While discovered over 40 years ago the field still lacks specific S-acylation inhibitors thus the potential benefit of pharmacologically targeting S-acyltransferases for human disease is still unknown. Here we report the identification of an orally bioavailable acyltransferase inhibitor SD-066-4 that inhibits the acyltransferase ZDHHC20. We identified a specific alanine residue that accommodates the methyl group of SD-066-4, thus providing isoform selectivity. SD-066-4 stably reduces EGFR S-acylation in Kras mutant cells and blocks the growth of Kras mutant lung tumors extending overall survival. We find that lung cancer patients harboring deletions in ZDHHC20 or ZDHHC14 concurrent with Kras alterations have a significant survival benefit, underscoring the translational importance of these enzymes.

## Main Text

ZDHHC20 is a member of a family of twenty-three protein-acyltransferases (PATs) containing the DHHC motif that catalyzes the reversible addition of fatty acids (predominately palmitate) to cysteine residues on protein substrates (*1, 2*). DHHC proteins are multi-pass integral membrane enzymes making them challenging to purify in their native state. The first crystal structure of a DHHC enzyme was recently solved for ZDHHC20 using a lipid cubic phase medium and confirmed the role of the lipid bilayer to the native structure of the enzyme and the binding pocket for acyl-CoA (*3*). Although DHHC proteins were identified as the mediator of cysteine acylation in 2002, there are still no specific drug-like inhibitors to any DHHC enzyme. The most widely used inhibitor, 2-bromopalmitate, covalently binds the hydrophobic pocket conserved between all DHHC enzymes, limiting its use as both a tool to study S-acylation and a drug (*4*). The recently developed inhibitor, cyano-myracrylamide (CMA) is more specific than 2-bromopalmitate but lacks enzyme selectivity (*5*). These challenges have hindered progress in understanding the regulation and function of S-acylation compared to other post-translational modifications.

ZDHHC20 is required for the initiation and progression of several cancer types in diverse models of tumorigenesis implicating it as a bon-a-fide target to inhibit tumorigenic processes. Collectively, deletion of ZDHHC20 prevents tumor initiation in a KrasG12D; p53 deficient genetically engineered mouse model (GEMM) of lung adenocarcinoma (LUAD), inhibits hepatocellular carcinoma tumor growth in vivo, and reduces growth of pancreatic cancer at metastatic sites (*6-8*). Mechanistically, S-acylation of EGFR on the C-terminal tail by DHHC20 is necessary for signaling through the PI3K-AKT axis, and as a result loss of EGFR S-acylation decreases PI3K signaling and Myc expression in Kras mutant LUAD cells [*6*]. Forced expression of an S-acylation resistant EGFR mutant in the Kras driven GEMM blocks lung tumor formation similar to ZDHHC20 ablation, confirming the mechanistic contribution of unmodified EGFR [*6*]. The potential value of ZDHHC20 as a drug target is further supported by the finding that the full body knock-out mouse of ZDHHC20 [IMPC MGI:1923215] is homozygous viable indicating this protein is not necessary for development and may only be essential for cancer cells.

### DHHC loss correlates with improved survival of Kras mutant NSCLC patients

Non-small cell lung cancers (NSCLC) account for 15% of all cancer-related deaths in the United States. To assess whether our observed protection against Kras driven NSCLC in the GEMM was applicable to human NSCLC, we examined the correlation between ZDHCC20 deletion in either KRAS mutant or KRAS wild-type lung adenocarcinoma patient data from The Cancer Genome Atlas (*9, 10*). Homozygous deletions in ZDHHC20 are found in 2.4% of patients (n=13) and only two of these patients harbor activating mutations or amplifications in KRAS, prohibiting survival analysis. Upon further examination of the TCGA database, we found homozygous deletions of ZDHHC14 in 2.3% of LUAD (n=12), where presence of either homozygous deletion (n=24) conferred a significant survival benefit in KRAS mutant lung cancer patients (Fig. 1A, fig. S1). Evidence of a protective function of ZDHHC20 loss in human patients, specifically those harboring activating KRAS alterations, aligns with our in vivo mouse studies that demonstrate ZDHHC20 loss inhibits KRAS driven tumorigenesis and furthered our interest in developing small molecule inhibitors of the enzyme.

**Figure 1.**
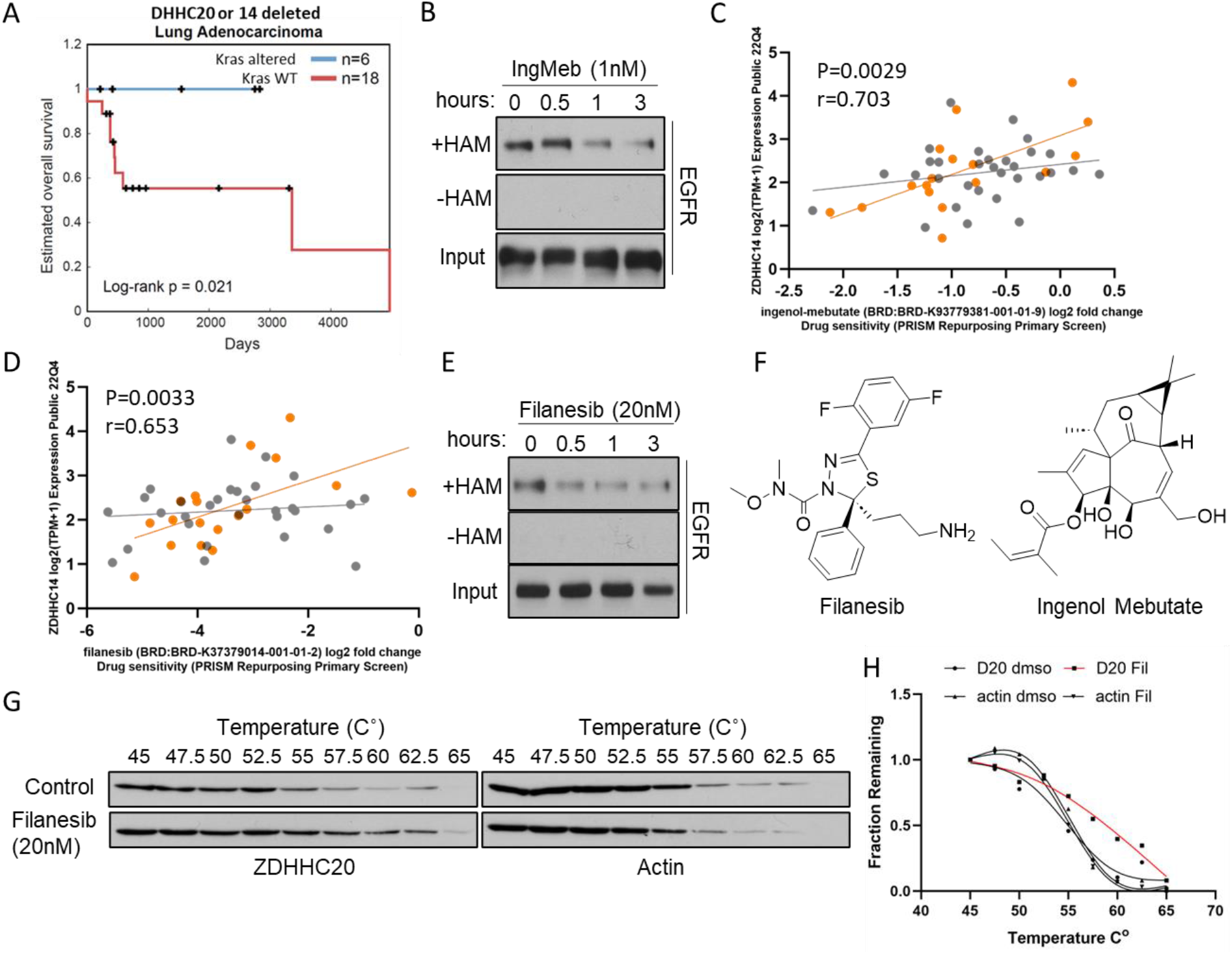
(**A**) Lung adenocarcinoma patients with deletions in ZDHHC20 or ZDHHC14. When paired with Kras alterations (blue) patients have a significant increase in overall survival compared to patients with wild type Kras (red) (TCGA database) Log-rank Test 0.021. (**B**) IngMeb inhibits EGFR S-acylation measured by ABE assay. Hydroxylamine is omitted as a negative control (-HAM) (**C**) Correlation between growth effect of IngMeb and ZDHHC14 mRNA expression in Kras mutant (orange) vs Kras wild type (gray) LUAD cell lines. DepMap cancer dependency database. (**D**) Correlation between growth effect of filanesib and ZDHHC14 mRNA in Kras mutant (orange) vs Kras wild type (gray) LUAD cell lines. (**E**) Treatment of H1975 cells with filanesib inhibits EGFR S-acylation. (**F**) Structures of filanesib and IngMeb. (**G, H**) Treatment of NCI-H23 cells expressing V5-tagged ZDHHC20 with filanesib increases the thermal stability of ZDHHC20-V5 by cellular thermal shift assay (CeTSA).

### Repurposing drugs to inhibit EGFR S-acylation

Considering the challenges related to the unique properties of ZDHHC20, we reasoned identifying a specific inhibitor would require assays in intact cells to capture the protein in its native state. A previous chemo-proteomic screen of proteins that bound the drug ingenol-mebutate (IngMeb) in live cells identified ZDHHC13 as a potential protein target, but the candidate was not studied further (*11*). IngMeb is a natural product from the plant Euphorbia peplus that is used topically to treat actinic keratosis, a precancerous skin condition (11). We found that 1nM IngMeb inhibited EGFR S-acylation within one hour of treatment (Fig. 1B). Based on our TCGA analysis of LUAD patient data, we reasoned that ZDHHC14 might be partially redundant with ZDHHC20 and that Kras mutant cell lines with low ZDHHC14 levels may be more sensitive to IngMeb. Using the DepMap database we found that the sensitivity of LUAD cell lines to IngMeb correlated negatively with ZDHHC14 mRNA expression levels, and this correlation was exclusive to Kras mutant LUAD cell lines (Fig. 1C) (*12*). We then searched DepMap for compounds with a similar correlation in Kras mutant LUAD cell lines and identified the inhibitor of the kinesis spindle protein (KSP), filanesib leading us to hypothesize that filanesib may also inhibit ZDHHC20 (Fig. 1D) (*13*). Treatment of cells with nanomolar concentrations of filanesib inhibited EGFR S-acylation, validating our methodology for identifying additional S-acylation inhibitors (Fig. 1E). For both IngMeb and filanesib there was no statistically significant correlation between drug effect and ZDHHC14 expression in Kras wild type cell lines (Fig. 1C, D). Unlike the natural product IngMeb, filanesib is a synthetic molecule with a readily accessible chemical structure (Fig. 1F), thus providing a promising starting point for further optimization. To determine if filanesib directly binds ZDHHC20, we used the cellular thermal shift assay (CeTSA), reasoning that the CeTSA would ensure we captured the relevant conformation of ZDHHC20 in live cells (*14*). The CeTSA revealed treatment of ZDHHC20-V5 expressing cells with filanesib increased the Tm of ZDHHC20-V5 compared to cells treated with the vehicle control (DMSO) (Fig. 1G, H). Filanesib treatment did not change the Tm of the negative control actin (Fig. 1G, H).

### Development of an S-acyl transferase inhibitor

To build on our chemogenomic approach, we used filanesib as a template to find structurally similar compounds that could be optimized to be selective for DHHC enzymes of interest, while avoiding KSP inhibition. Using Rapid Overlay of Chemicals Shape (ROCS) computational methods (*15, 16*), we searched the NIH Molecular Libraries Small Molecule Repository of approximately 350K compounds for molecules with no known biological function that share 3D structural similarities with filanesib (Fig. 2A). The pendant amine in filanesib (also present in KSP inhibitor ispinesib) is critical to its activity as an inhibitor of KSP and was therefore excluded from the shape searching query (*17*). This ligand-based screening approach resulted in three compounds that were purchased and tested experimentally in an acyl-biotinyl exchange assay to measure EGFR S-acylation in NCI-H1975 lung cancer cells (fig. S2A). One of these compounds, Compound-1, inhibits EGFR S-acylation at concentrations as low as 1μM after treatment for one hour, but has no effect on Ras S-acylation, a substrate of ZDHHC9, suggesting isoform selectivity (fig. S2B). These structurally related compounds begin to outline the structure-activity relationships (SAR) for inhibition of EGFR S-acylation. Building on this, we synthesized multiple compounds to probe the SAR of Compound-1, leading to the identification of the most potent inhibitor to date, SD-066-4 (Fig. 2B). Treatment of NCI-H1975 lung adenocarcinoma cells at low micromolar concentrations of SD-066-4 reduces EGFR S-acylation within one hour (Fig. 2B), while preserving spindle separation and Ras S-acylation and as a result was designated our lead compound for further characterization (Fig. 2C, fig. S3).

**Figure 2.**
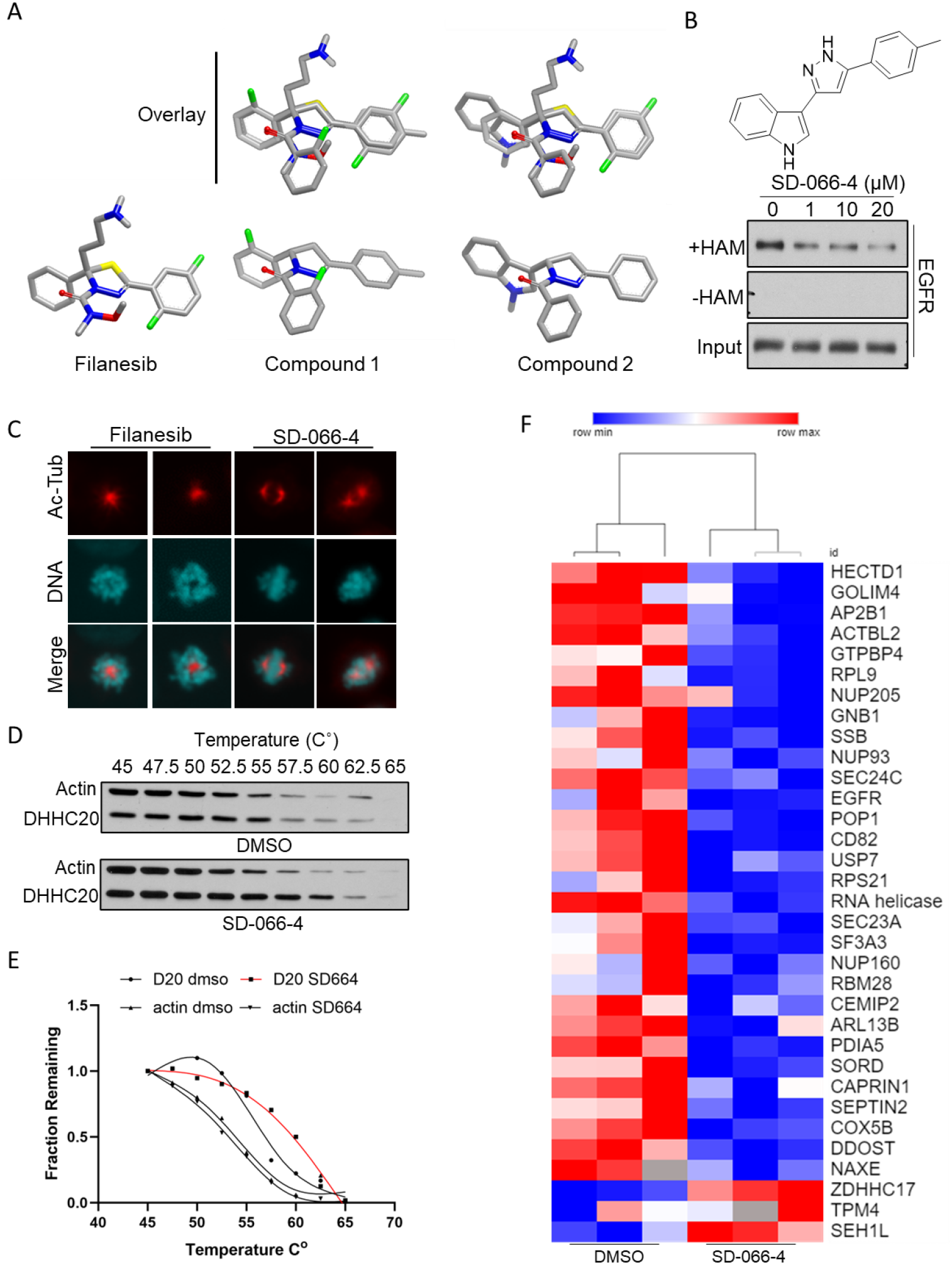
(**A**) Overlay of filanesib with ROCS identified compounds 1 and 2. (**B**) Treatment of H1975 cells with SD-066-4 for 1 hour decreases EGFR S-acylation. (**C**) Immunofluorescence image of centrosome separation during mitosis. Kinesin inhibitor filanesib (20nM) inhibits centrosome separation during mitosis. Centrosome separation is unimpaired in SD-066-4 (10μM) treated cells. Acetylated tubulin antibody labels the mitotic spindles. (**D**)Treatment of NCI-H23 cells expressing ZDHHC20-V5 with SD-066-4 increases the thermal stability of ZDHHC20-V5 measured by CeTSA. (**E**) Quantification of CeTSA immunoblot. (**F**) Changes in the S-acyl proteome upon treatment with SD-066-4.

The binding of SD-066-4 appears to be direct as the treatment of ZDHHC20-V5 expressing cells with SD-066-4 increased the Tm of ZDHHC20-V5 compared to vehicle control treatment (DMSO) (Fig. 2D, E). The Tm of the negative control actin did not change upon treatment with SD-066-4 (Fig. 2D, E).

To comprehensively examine the changes in protein S-acylation induced by SD-066-4, we performed proteomics by LC-MS/MS on ABE isolated S-acylated proteins from NCI-H23 KrasG12C mutant LUAD cells treated with one micromolar SD-066-4 or DMSO for three hours. In total we identified 1,177 S-acylated proteins by three or more unique peptides and in cells treated with SD-066-4 EGFR S-acylation was reduced 1.7-fold. Using EGFR as a reference for a physiologically relevant decrease, we found that only 2.5% of the S-acylated proteome decreased upon SD-066-4 treatment by greater than 1.5-fold with a p-value of less than 0.05 (Fig. 2F, Table S1). These results indicate high selectivity for the S-acylation inhibitor in Kras mutant LUAD cells.

### Modeling the binding site of SD-066-4

Seeking to understand the molecular interaction between SD-066-4 and DHHC20, we performed docking studies (see methods). The resulting model suggests a binding pose between the lipid-binding pocket and the nucleotide-binding site (Fig. 3A). This pose is supported by the SAR of SD-066-4 in which we found that methylation of the indole abrogates activity since it would introduce a steric clash with the protein surface. The model also predicts ZDHHC14 should be resistant to SD-066-4 because of a steric clash introduced by an alanine to leucine variation between ZDHHC20 and ZDHHC14 (Fig. 2B). We confirmed this model by showing that NCI-H1975 cells overexpressing ZDHHC14 cloned from mouse (mZDHHC15) are resistant to SD-066-4 mediated inhibition of EGFR S-acylation (Fig. 2C). To test this model further, we developed an analog of SD-066-4 that lacks methylation of the benzyl group, SD-128 (Fig. 2D). We found that SD-128 induced a potent dose dependent inhibition of EGFR palmitoylation in cells overexpressing mZDHHC14-HA but had minimal effect on the parental cells without mZDHHC14-HA suggesting that the leucine in ZDHHC14 serves as a “gatekeeper” (Fig. 2E).

**Figure 3.**
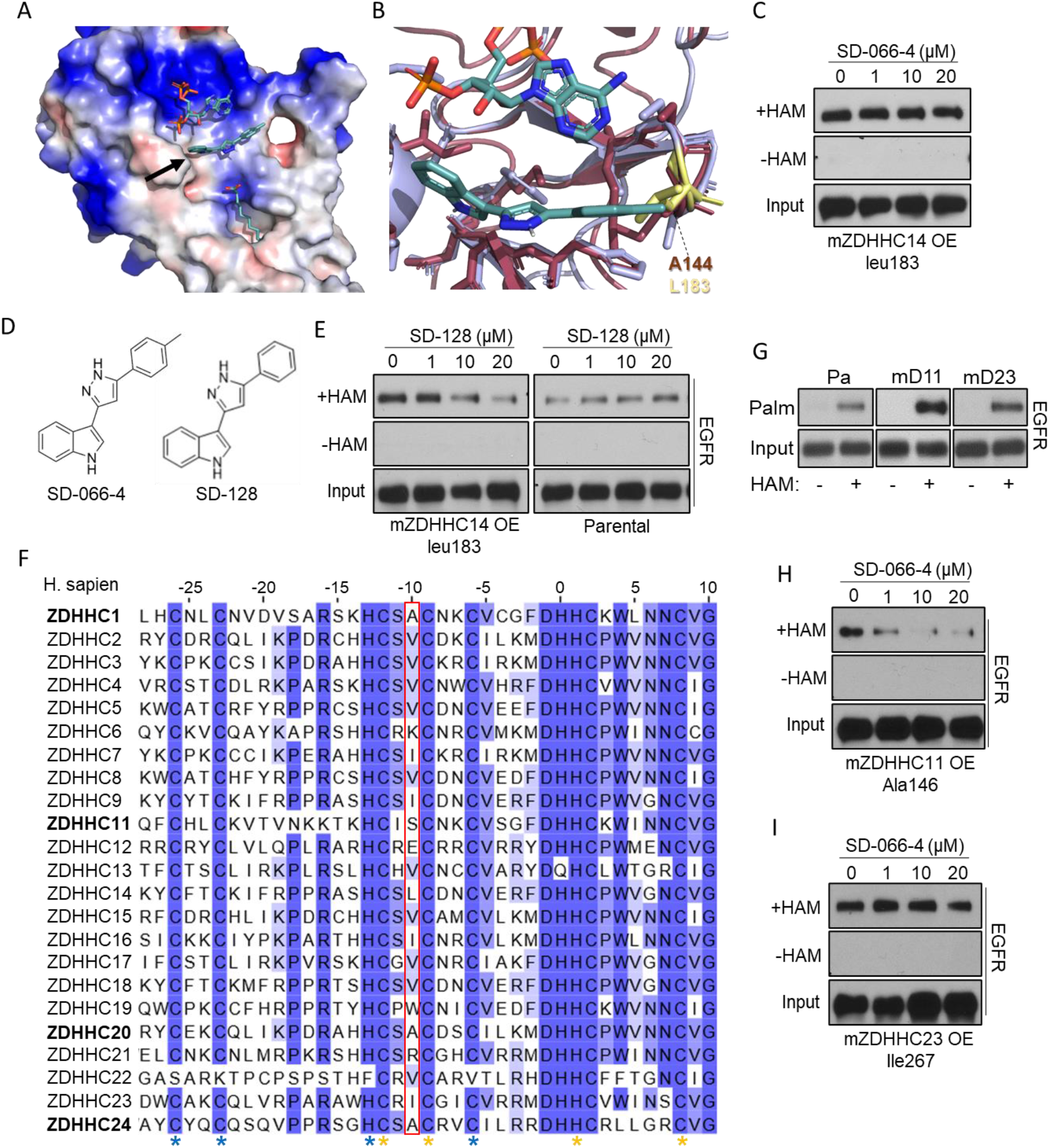
(**A**) Modeling of potential SD-066-4 binding site (arrow) on ZDHHC20. (**B**) Methyl group of SD-066-4 clashes with leucine (yellow) in ZDHHC14 (AlphaFold). (**C**) Cells expressing mZDHHC14 are resistant to SD-066-4 mediated inhibition of EGFR S-acylation. (**D**) Chemical structures of SD-066-4 compared to SD-128. (**E**) Cells expressing mZDHHC14 are permissive to SD-128 which has no effect on parental cells. (**F**) ZDHHC enzymes in bold have small amino acid side groups in position of alanine 144 (Red box) in ZDHHC20. Sequence conservation highlighted in blue (darker is more conserved). Asterisks indicate conserved CCHC zinc finger motifs. (**G**) Overexpression of mZDHHC11 or mZDHHC23 increases EGFR S-acylation. (**H, I**) EGFR S-acylation is inhibited by SD-066-4 in cells overexpressing mZDHHC11 but not in cells overexpressing mZDHHC23.

Analysis of the predicted structures of all twenty-three DHHC enzymes (AlphaFold protein structure database) reveals only ZDHHC1, 11, 20 and 24 contain a small side chain amino acid at the spatial position of alanine 144 within one of the conserved CCHC zinc finger motifs. The remaining nineteen DHHC enzymes contain bulky residues at this position, thus providing a rationale for the observed selectivity of SD-066-4 (Fig. 2F). To test our model for DHHC enzyme selectivity, we forced expression of either mZDHHC11 or mZDHHC23 in NCI-H1975 cells and assessed EGFR acylation and sensitivity to SD-066-4. While expression of either mZDHHC11 or mZDHHC23 increases EGFR acylation (Fig. 3G), only mZDHHC11, which contains an alanine in the same spatial position as alanine 144 in ZDHHC20 was responsive to SD-066-4 (Fig. 2H). mZDHHC23 contains an isoleucine at the alanine position and likely underlies the resistance to SD-066-4 at identical concentrations of cells overexpressing ZDHHC23 (Fig. 3I). The observed selectivity for ZDHHC20 and mZDHHC11 supports the model that alanine 144 provides the observed selectivity of SD-066-4 for ZDHHC20.

### Sensitivity of Kras mutant cells to SD-066-4

Our previous studies showed that genetic deletion of ZDHHC20 in Kras mutant lung cancer cells reduced levels of AKT activating phosphorylation and Myc protein, we therefore asked if treatment with SD-066-4 induced a similar effect. In Kras mutant cells pAKT and Myc levels initially increased but steadily decreased by 6 hours of treatment and were lower than control treated cells by twenty hours (Fig. 4A). In contrast the Kras wild-type cell line NCI-H1975 pAKT levels were constant over the time course while Myc also initially increased but returned to control treated levels by six hours (Fig. 4A). To determine if the kinetic difference in signaling correlates with EGFR S-acylation we measured EGFR S-acylation in SD-066-4 treated Kras mutant cells compared to Kras wild-type cells over twenty hours. In the KrasG12S mutant cell line A549 EGFR S-acylation was maximally inhibited after six hours of 1μM SD-066-4 treatment and remained inhibited after twenty hours (Fig. 4B). In contrast, Kras wild-type line NCI-H1975 was inhibited at one hour treatment but S-acylation recovered by six hours (Fig. 4B).

**Figure 4.**
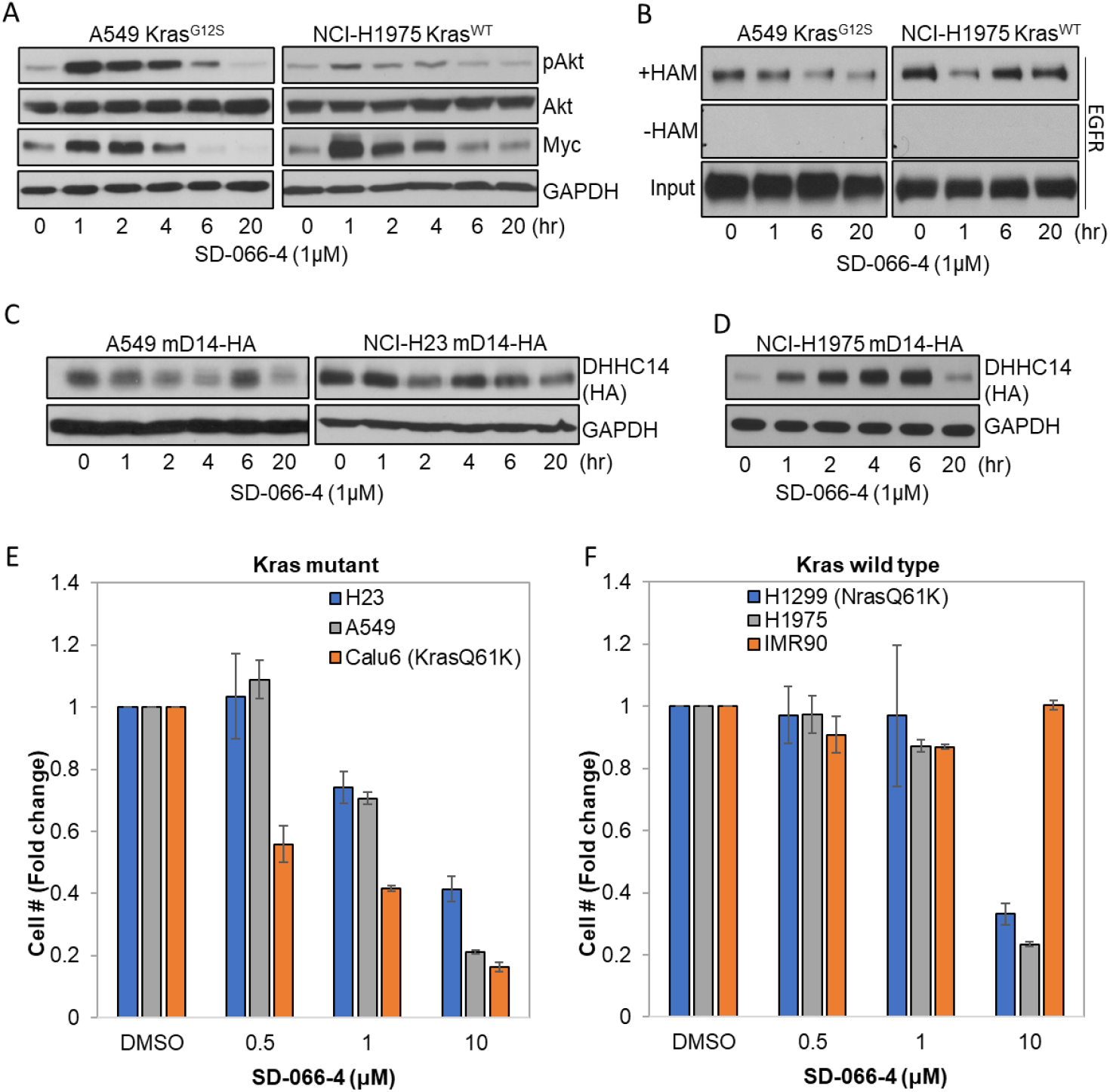
(**A**) Altered pAKT (T308) and Myc protein levels in Kras mutant and Kras wild type LUAD cell lines. (**B**) Inhibition of EGFR S-acylation over time in Kras mutant and Kras wild type LUAD cell lines. (**C, D**) Treatment of Kras mutant LUAD cells decreases ZDHHC14-HA protein levels over time (C), in contrast to Kras WT LUAD cells where ZDHHC14-HA increases (D). (**E, F**) Cell proliferation of Kras mutant and Kras wild-type LUAD cell lines and normal lung fibroblast cells line with dosage increase in SD-066-4.

We hypothesized that the recovery in EGFR S-acylation observed in Kras wild-type cells after six hours of SD-066-4 treatment may be caused by increased expression of an alternate transferase with overlapping function. To test this hypothesis, we established A549 KrasG12S, NCI-H23 KrasG12C and H1975 Kras wild-type cell lines expressing HA tagged ZDHHC14 and measured ZDHHC14 protein levels over twenty hours of SD-066-4 treatment. ZDHHC14-HA levels decreased modestly after twenty hours in A549 and NCI-H23 (Fig. 4C). In contrast, ZDHHC14-HA protein levels rapidly increase by one hour of treatment in NCI-H1975 cells and peaked at six hours followed by a decrease at twenty hours (Fig. 4D).

Lastly, we compared the sensitivity to ZDHHC20 inhibition by SD-066-4 of Kras mutant and Kras wild-type lung cancer cells. We found that proliferation of the Kras mutant lung cancer lines NCI-H23, A549 and Calu6 were inhibited with an IC50 of approximately 2μM SD-066-4 whereas the Kras wild-type lung cancer cell lines NCI-H1975 and NCI-H1299 required 5-10 μM SD-066-4 to achieve similar growth inhibition (Fig. 4E, F). Inhibition of proliferation was specific to cancer cell lines as SD-066-4 was unable to inhibit proliferation of the lung fibroblast cell line IMR90 at identical concentrations and indicates low nonspecific toxicity (Fig. 4F).

### SD-066-4 inhibits lung cancer tumor growth

Initial assessment of the pharmacokinetic properties of the SD-066-4 revealed acceptable pharmacological properties: 10 mg/kg IP or PO dosing followed by plasma level quantitation over the following 24 hours revealed maximum concentrations of 1270 ng/ml and 219 ng/ml and half-lives of 1.9 and 4.6 hours, respectively (Fig. 5A). Based on these favorable pharmacokinetic properties, we assessed the therapeutic potential of SD-066-4 in preclinical xenograft models.

**Figure 5.**
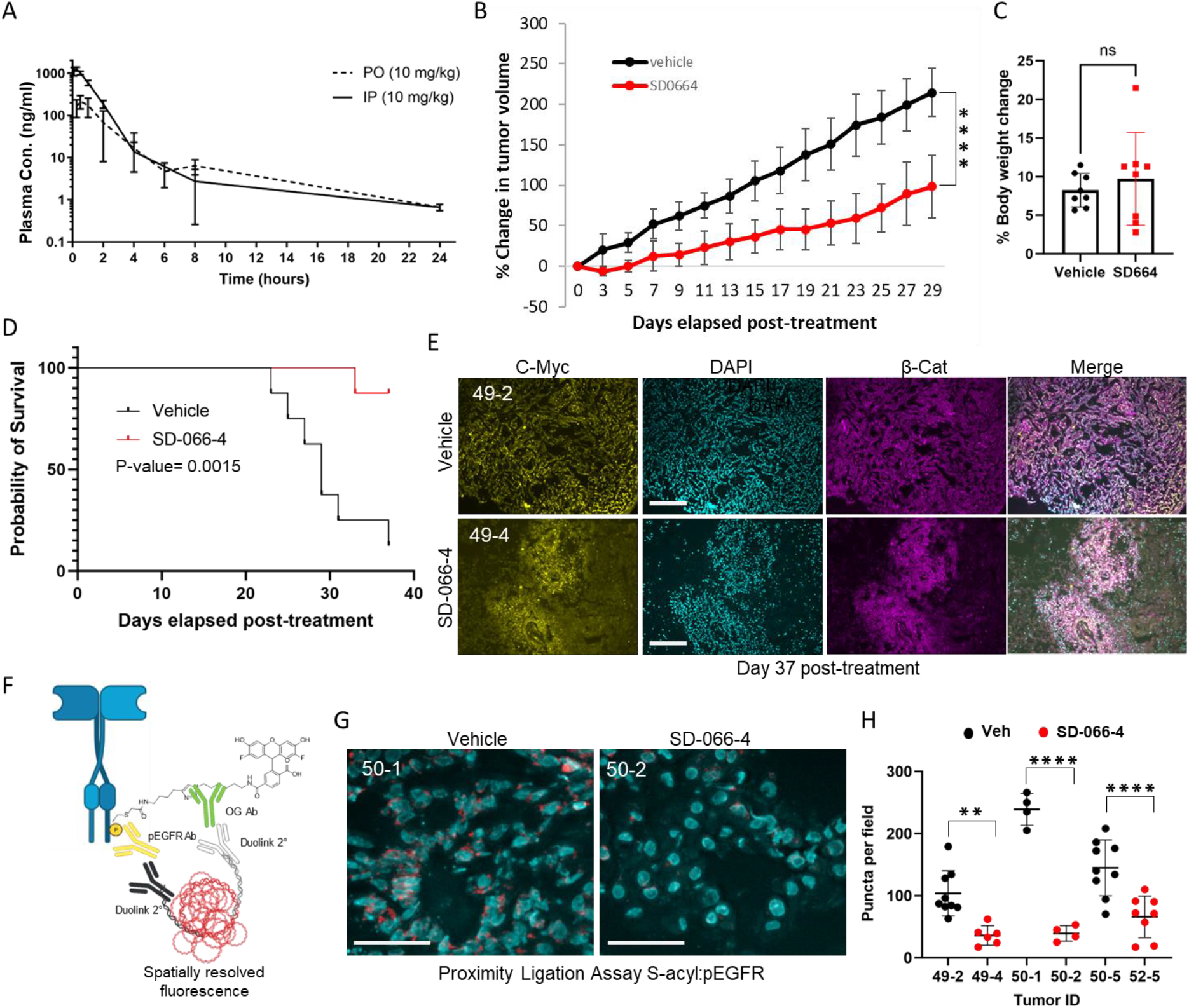
(**A**) Murine PK of SD-066-4 dissolved in PEG400 and mixed 1:1 with 20% captisol delivered per oral (PO) and intraperitoneal (IP). (**B**) Percent tumor volume change of vehicle vs. SD-066-4 treated tumors over 29 days. (**C**) Percent body weight change of vehicle or SD-066-4 treated animals at endpoint. (**D**) Kaplan Myer Plot of probability of survival of vehicle vs. SD-066-4 animals. Log-rank P value=0.0015. (**E**) Immunofluorescence staining of frozen tumor sections probed with anti-Myc (yellow) anti-β-catenin (magenta) and DAPI (cyan). 40X magnification, scale bar = 250μm. (**F**) Schematic of S-acyl PLA to detect S-acylated phosphorylated EGFR. (**G**) Representative image of frozen tumor section treated and analyzed with S-acyl PLA from vehicle and SD-066-4 treated tumors. Scale bar = 45μm. (**H**) Quantification of PLA puncta of vehicle and SD-066-4 treated tumors counted from deconvolved z-series. **P=0.0018, ****P<0.0001; 2way ANOVA.

The KrasG12S lung cancer cell line A549 was engrafted in the flanks of NOD SCID mice and when tumors reached a volume of 100mm3 mice were treated with SD-066-4 or vehicle control by oral gavage to find an optimal dosing regimen. While daily treatment with 50mg/kg with SD-066-4 had no effect on tumor growth, increasing the dose to 100mg/kg daily showed growth inhibition over seven days (fig. S5A, B). The study was repeated and when tumor volume reached 300mm3 mice were treated daily with 100mg/kg by oral gavage. After five days of treatment the average percent change in volume was 0% compared to 29.2% for the vehicle control (fig. S5C). The first animal in the vehicle cohort reached the endpoint tumor volume (1000mm3) on treatment day twenty-three and the average change for vehicle treated tumors was 174.5% compared to 59.0% for SD-066-4 treated tumors (Fig. 5B; fig. S5D). There was no significant difference in percent body weight change between vehicle and SD-066-4 treated animals after day thirty-seven, indicating there is no overt general toxicity at the therapeutic dose (Fig. 5C). Treatment with SD-066-4 as a single agent increased overall survival with seven of the eight control treated animals reaching end point tumor volume by the end of the study compared to only one of the eight SD-066-4 treated animals (Fig. 5D).

To molecularly characterize the effect of SD-066-4 in vivo, tumors were sectioned and analyzed by immunofluorescence staining. The tumors that showed the greatest growth inhibition contained restricted regions expressing the cell membrane markers β-catenin or E-cadherin and c-Myc protein and large areas with sparse nuclei and low levels of the membrane markers and nuclear c-Myc. In contrast the vehicle control tumor was mostly intact with uniform expression of both Myc and β-catenin/E-cadherin across the tumor section (Fig. 5E; fig. S6A, B). Since the most effectively inhibited tumors contained only small regions of intact healthy tumor tissue, we reasoned the conventional ABE assay on tumor lysate would not provide accurate levels of EGFR S-acylation in the tumor cells, and we instead used a recently developed method to spatially resolve S-acylated EGFR in situ (*18*). The method involves applying a variation of the S-acyl exchange assay on frozen tumor sections to label S-acylated cystine residues with the fluorophore Oregon Green and detecting the dually S-acylated and phosphorylated EGFR by a Duolink proximity ligation assay with antibodies to Oregon Green and phosphorylated EGFR which generates fluorescent puncta (Fig. 5F, G; fig. S7A). The PLA puncta per field were quantified revealing a reduction of S-acylated pEGFR in SD-066-4 treated animals compared to the vehicle control tumors (Fig. 5H). As previously reported, we found inhibition of ZDHHC20 increases EGFR phosphorylation at tyrosine 1068 by immunofluorescence staining, confirming the reduction in PLA puncta was due to decreased EGFR S-acylation not decreased phosphorylation (fig. S7B) (*19*).

These studies provide a drug-like small molecule inhibitor, SD-066-4, of ZDHHC20, demonstrate its potential for therapeutic use in Kras mutant LUAD under oral administration and provide a roadmap for inhibition of other members of this enzyme class. Crucially, these studies begin to identify key residues that impart drug selectivity for S-acylation inhibitors, revealing possibilities for drugging additional members of the ZDHHC family using altered versions of SD-066-4. The structures of the DHHC enzymes are highly conserved among the family members with the deep hydrophobic lipid-binding cavity being the most obvious site for small molecule inhibitor design. The recently developed DHHC inhibitor CMA was designed around a saturated alkane moiety that targets the lipid-binding pocket and is therefore not selective for individual or subsets of DHHC enzymes. Recently, Artemisinin was also shown to be a palmitoyl transferase inhibitor, but the selectivity is unknown, and the complex sesquiterpene lactone structure makes optimization challenging (*20*). Our data suggests that SD-066-4 binds outside the lipid binding pocket and while the affinity may be reduced compared to lipid pocket binding compounds, SD-066-4 displays a unique selectivity profile. This can be explained by the identification of key residues that impart drug selectivity for SD-066-4 and provide insight for drugging additional members of the ZDHHC family using similar molecules.

Biologically, the development of selective small molecules inhibitors allowed us to acutely inhibit S-acylation, revealing previously unknown dynamic changes in oncogenic signaling networks. This led to the discovery that cells need to maintain EGFR S-acylation in a Kras mutant setting. This observation is further supported by our finding that EGFR expressing cells respond to the introduction of oncogenic Kras by increasing EGFR S-acylation by greater than tenfold. This adaptive response is consistent with the requirement for ZDHHC20 mediated S-acylation to optimize growth signaling in the presence of oncogenic Kras and provides a molecular mechanism for the mutual exclusivity of Kras and EGFR activation.

Furthermore, we demonstrate the therapeutic potential of an orally administered drug-like small molecule inhibitor of ZDHHC20 to treat KRASmut lung adenocarcinoma, thus providing an alternative therapeutic strategy to recently developed covalent Kras inhibitors that are being challenged in the clinic with rapid evolution of drug resistance and which only inhibit subsets of the Kras mutants. We envision the ability to drug ZDHHCs will open new avenues for drug discovery for a plethora of diseases including lung cancer, leukemia, Friedreich’s ataxia and malaria (*21-23*).

## Acknowledgments

The cDNAs for mouse ZDHHC11, 14, and 23 were generous gifts from Dr. Masaki Fukata. We are grateful to OpenEye Scientific for an Academic License to support our research.

## Author contributions

Conceptualization: ESW, GMB

Methodology: ESW, GMB, DCB, NA

Investigation: JYL, MA, AM, TM, SD

Project administration: ESW, GMB

Supervision: ESW, GMB, DCB

Writing – original draft: ESW, GMB

Writing – review & editing: ESW, GMB, DCB, NA

## Data and materials availability

All source data are provided with this paper or the supplementary materials. Materials are available upon reasonable request and may require a Material Transfer Agreements (MTAs).

## Supplementary Materials

Materials and Methods

Figs. S1 to S7

Tables S1

Data S1

