## Supplemental Information for "A selective S-acyltransferase inhibitor suppresses tumor growth"

**The PDF file includes:**

Materials and Methods
Figs. S1 to S6
Tables S1

**Other Supplementary Materials for this manuscript include the following:**

Data S1

### Materials and Methods

Cell culture

A549, Calu6, NCI-H1975 and IMR90 cells (ATCC) were cultured in DMEM containing 10% FBS. NCI-H23, NCI-H1299 were maintained in RPMI supplemented with 10% FBS. Cell lines tested for mycoplasma contamination using MycoAlert Mycoplasma Detection Kit (Lonza). All cell lines were free of contaminants.

Antibodies

Anti-DHHC20 (HPA014702) antibody was purchased from Sigma-Aldrich. Anti-EGFR-XP (catalog no. 4267S), pERK (catalog no. 4370S), ERK (catalog no. 4695S), pS473-AKT (catalog no. 4060S), pT308-AKT (catalog no. 13038S), AKT (catalog no. 4691S), Ras (catalog no. 3339S), β-catenin (catalog no. 8480S), Anti-Myc (catalog no. 2272S) and β-actin (catalog no. 4970S) were obtained from Cell Signaling Technologies. Anti-acetylated tubulin was purchased from Santa Cruz. Phosphorylated EGFR (Y1068) (Novus Biologicals #MAB3570) and for fluorescein/OG 488 (Life Technologies A-889)

Small molecule compounds

Candidate molecules identified by ROCS analysis were purchased from MCULE Inc. (Palo Alto, CA). Compound-1 (MCULE-5940587479); Compound-2 (MCULE-2264298670); Compound-3 (MCULE-4656029049). Filanesib was purchased from MedChemExpress (HY-15187). Ingenol Mebutate was purchased from Cayman Chemical (#16207).

Synthesis of Novel Molecules

Solvents were removed under reduced pressure using an IKA rotary evaporator equipped with an IKA MVP10 basic vacuum pump. Dry solvents and reagents were purchased from commercial suppliers and used without further purification. Thin-layer chromatography was performed on Merck silica gel 60F254 plates. Flash column chromatography was conducted using a Teledyne ISCO CombiFlash NextGen 300+ purification system using 4g, 12g, or 24g RediSep Rf silica cartridges. Thermo Scientific silica gel (35-70 μm) was used for manual chromatography columns.

Analytical LC-MS was performed using a system comprised of an Agilent 1260 Infinity II HPLC instrument equipped with an Agilent InfinityLab LS/MSD XT MS detector with electrospray ionisation. The system ran with a positive and negative switching mode and UV diode array detector using an Agilent ZORBAX SB-C18 RRHT (50 mm × 4.6 mm × 1.8 μm) column and gradient elution with two binary solvent systems: MeCN/$H_2O$ or MeCN/$H_2O$ plus 0.1% formic acid. Preparative HPLC was performed using a system comprising an Agilent 1260 Infinity II HPLC system equipped with an Agilent 1290 Prep Bin Pump, an Agilent Prep-C18 column (250 mm × 21.2 mm × 10 μm), and an Agilent prep autosampler and fraction collector. The system ran using a UV diode array detector and purification was performed using a gradient elution using a MeCN/$H_2O$ binary solvent system.

NMR analysis was conducted using a Bruker NEO400 spectrometer equipped with a 5mm BBFO IProbe ($^1$H = 400 MHz and $^{13}$C = 100 MHz) or a Bruker NEO600 spectrometer equipped with a Prodigy BBO cryoprobe ($^1$H = 600 MHz and $^{13}$C = 150 MHz). Chemical shifts are quoted in parts per million (ppm) and coupling constants are given in Hz. Splitting patterns have been abbreviated as follows: s (singlet), d (doublet), dd (doublet of doublets), ddd (doublet of doublets of doublets), td (triplet of doublets), t (triplet), q (quartet) and m (multiplet). NMR data is reported in the format: ppm (number of protons, splitting pattern, coupling constant).

Synthetic Scheme:

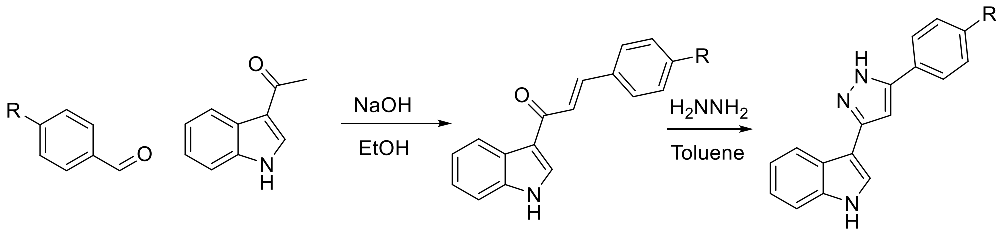


General Procedure A: Claisen Schmidt Condensation

To a solution of a benzaldehyde (1 eq.) in ethanol (2 ml per mmol aldehyde) was added aqueous sodium hydroxide (3M, 1 ml of mmol of aldehyde) followed by 3-acetylindole (1 eq.) and the reaction was stirred at r.t. for 2 hours. The resulting yellow precipitate was collected by filtration, washed with cold water (3 x 10 ml). The product was purified by recrystallization from methanol.

General Procedure B: Pyrazole Formation

To a solution of (E)-1-(1H-indol-3-yl)-3-arylprop-2-en-1-one (1 eq.) in toluene (1 ml per gram) was added hydrazine hydrate (2.5 eq.) dropwise and the reaction was heated to reflux for 8 hours. The reaction mixture was then concentrated in vacuo, resuspended in water, adjusted to pH 7.0 and extracted with ethyl acetate (3 x 15 ml). The combined organic layers were washed with water and brine, dried over MgSO4 and concentrated in vacuo. The resulting residue was purified by column chromatography eluting with 30-75% ethyl acetate in hexane.

SD-066-4 3-(5-(*p*-tolyl)-1H-pyrazol-3-yl)-1H-indole

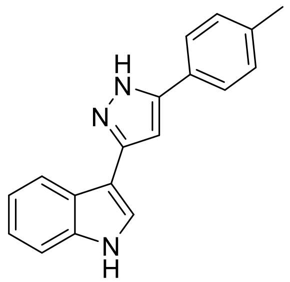


3-(5-(*p*-tolyl)-1H-pyrazol-3-yl)-1H-indole (SD066-4) was synthesized following general procedures A and B from tolualdehyde and 3-acetyl indole.
$^1$H NMR (400 MHz, $CDCl_3$) δ 9.14 (s, 1H), 8.34 (dd, *J* = 6.5, 2.8 Hz, 1H), 7.35 (dt, *J* = 7.5, 3.1 Hz, 1H), 7.31 – 7.28 (m, 2H), 7.24 (d, *J* = 2.8 Hz, 1H), 7.17 – 7.06 (m, 4H), 5.52 (dd, *J* = 11.5, 4.2 Hz, 1H), 3.75 (dd, *J* = 17.0, 11.5 Hz, 1H), 3.12 (dd, *J* = 17.0, 4.2 Hz, 1H), 2.53 (s, 3H)
$^{13}$C NMR (101 MHz, CDCl3) δ 168.6, 151.3, 139.2, 137.2, 136.8, 125.6, 124.7, 123.5, 122.2, 121.5, 111.6, 109.7, 58.5, 43.5, 22.2, 21.1.
LC/MS: Calculated for $C_{18}H_{16}N_3$ $[M+H]^+$: 274.1; Found: 273.9.

SD-128 3-(5-phenyl-1H-pyrazol-3-yl)-1H-indole

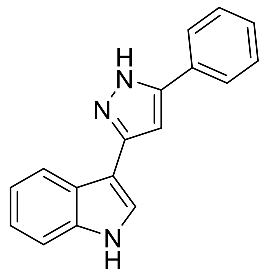


3-(5-phenyl-1H-pyrazol-3-yl)-1H-indole (SD-128) was synthesized following general procedures A and B from benzaldehyde and 3-acetyl indole.
$^{1}$H NMR (400 MHz, MeOD-d4) δ 7.95 (dd, *J* = 6.9, 1.8 Hz, 1H), 7.85 (d, *J* = 7.1 Hz, 1H), 7.70 (s, 1H), 7.48 – 7.43 (m, 4H), 7.39 – 7.34 (m, 2H), 7.19 (pd, *J* = 7.1, 1.5 Hz, 4H), 6.99 (s, 1H).
$^{13}$C NMR (101 MHz, MeOD) δ 138.3, 133.1, 129.9, 129.4, 126.9, 126.5, 124.5, 123.2, 121.2, 120.5, 112.7, 107.4, 100.8.
LC/MS: Calculated for $C_{17}H_{13}N_3$ $[M+H]^+$: 260.1; Found: 260.1.

Plasmids and Generation of Stable Cell Lines

Human ZDHHC20 and ZDHHC14 from pLV-tetO with an HA tag (Addgene plasmid #19763) (20) was gateway cloned into the pLX304 backbone with a blasticidin resistance marker and V5 tag (Addgene). Lentivirus of pLX304-ZDHHC20/14 was generated using HEK293T cells with Gag, VSVG and Rev plasmids using TransIT-LT1 (Mirus) according to manufacturer's instructions. H23 cells that were transduced with lentivirus and selected with blasticidin (10 µg/ml) together for several passages. Inducible cell lines, wildtype EGFR KRAS4B(WT) or mutant KRAS4B(G12V) in pLenti-PGK-hygromycin resistance with an HA tag (Addgene plasmid #35633) (20) was generated using HEK293T cells with Gag, VSVG and Rev plasmids using TransIT-LT1 (Mirus) according to manufacturer's instructions. NIH3T3 cells infected with pTRIPZ constructs and selected with puromycin were subsequently infected with KRAS4B(G12V)-HA are described in Kharbanda 2020.

TCGA analysis

LUAD TCGA copy number and somatic mutation data was downloaded through Xena browser1. Gene-level copy number values were estimated using GISTIC22 method where converted values -2,-1,0,1,2, represent homozygous deletion, single copy deletion, diploid normal copy, low-level copy number amplification, or high-level copy number amplification. For ZDHHC20 and ZDHHC14 homozygous deletions were considered as deleted samples. For Kras, point mutation (G12 substitutions) and high-level copy number amplifications were considered mutated. The influence of ZDHHC20 and ZDHHC14 deletions over patient outcomes in presence of Kras mutations was evaluated by comparing overall (OS) and disease stable survival (DSS)3 in ZDHHC20 or ZDHHC14 deleted patients with WT vs mutated Kras using log-rank test.

Docking studies

The crystal structure of ZDHHC20 (PDB ID:6BML) was prepared for docking but no constraints were enforced on the site of binding. Low energy conformers of SD-066-4 were generated using Omega2 and docked into ZDHHC20 using FRED.

Acyl-biotinyl exchange (ABE) assay

ABE assay was performed as described in Runkle 2016.

Immunoblot analysis

Cell lysates were prepared in 1% Triton-X-100 buffer, including Tris-HCl (pH 7.5) and sodium chloride solution (NaCl). Lysates were analyzed by immunoblotting with the following antibodies: Primary antibodies were diluted 1:1000 in 5% bovine serum albumin (BSA) dissolved in tris buffered saline plus 0.01% tween (TBST) and incubated overnight at 4ºC. Immune complexes were detected with horseradish peroxidase-conjugated secondary antibodies diluted in 5% non-fat milk dissolved in TBST and enhanced chemiluminescence (ECL) (Thermo Scientific).

Cellular Thermal Shift Assay

NCI-H23 cells expressing ZDHHC20-V5 were treated with Filanesib (20nM) or SD-066-4 (20µM) or DMSO for one hour then washed with PBS and harvested by scraping and distributed to PCR tubes in PBS containing drug or vehicle DMSO. The remaining steps of CeTSA was performed as described in Reinhard et al. 2015.

Sample preparation for S-acyl proteomic analysis

Proteomic samples were prepared in triplicate (1 x 60mm TC plate per replicate) by the ABE assay as described previously. For bottom-up proteomics analysis equal volumes of ABE processed streptavidin agarose beads (30µl of beads per sample) were taken across samples and washed in 50mM triethyl ammonium bicarbonate buffer (TEABC). Samples were spiked with 50ng of BSA to be used for normalization. Samples were reduced using 5 mM DTT (Dithiothreitol) at 60 degrees for 30 minutes followed by alkylation using 20mM IAA (Iodoacetamide) for 15 minutes in the dark. Beads were diluted by adding 50ul of 50 mM TEABC Ph 8.0, and trypsinized using 500 ng of modified sequence-grade trypsin (Promega, Madison, WI) at 37 degrees overnight.

Peptide desalting was carried out with the C-18 StageTip method. The C-18 material was stacked onto 200 µl tips, activated with 100% ACN, and equilibrated with 0.1% formic acid. Peptide samples were resuspended in 0.1% formic acid and loaded onto the C-18 StageTip. The sample was passed two times, followed by washing with 0.1% formic acid and elution with 40% ACN in 0.1% formic acid. The dried fractions were stored at -20°C until LC-MS/MS analysis.

LC-MS/MS analysis

The C18 cleaned peptides were analyzed on Thermoscientific Orbitrap Exploris 240 mass spectrometer interfaced with Thermoscientific UltiMate 3000 HPLC and UHPLC Systems. Peptide digests were reconstituted in 0.1 % formic acid and separated on an analytical column (75 µm × 15 cm) at a flow rate of 300 nL/min using a gradient of 1–25 % solvent B (0.1 % formic acid in 100 % acetonitrile) for the first 100 minutes and 25–30 % for next 5 minutes, 30-70 % for 5 minutes, 70–1 % for next 5 minutes. The total run time was set to 120 min. The mass spectrometer was operated in data-dependent acquisition mode. A survey full scan MS (from m/z

400–1600) was acquired with a resolution of 6000 Normalized AGC target 300. Data were acquired in topN with 20 dependent scans. Fragmentation was carried out using normalized collision energy of 37 % and detected at a mass resolution of 1500. Dynamic exclusion was set for 8 s with a 10-ppm mass window.

Proteomic data analysis

MS/MS searches were carried out using SEQUEST search algorithms against a Uniprot database for humans using Proteome Discoverer (Version 3.0, Thermofisher Scientific, Bremen, Germany). The workflow included Spectrum files, Spectrum selector, SEQUEST search nodes, Target decoy PSM validator, peptide validator, event detector, precursor quantifier. Data was searched in label free quantification mode using unique peptides for quantification. Oxidation of methionine, N-terminal protein acetylation, N-ethylmaleimide of cysteine, deamidation of asparagine, glutamine, and phosphorylation of serine, threonine and tyrosine were used as dynamic modifications and carbamidomethylation of cysteine was set as static modification. MS and MS/MS mass tolerances were set to 10 ppm and 0.05 Da, respectively. Trypsin was specified as protease and a maximum of two missed cleavage was allowed. Target-decoy database searches were used for calculation of false discovery rate (FDR) and for peptide identification FDR was set at 1%. Feature mapper and precursor ion quantifier were used for label-free quantification. Normalization was set as specific protein amount and BSA was used for normalization.

NSCLC Xenografts

Four to six-week-old SCID beige mice (CB17.Cg-PrkdcscidLystbg-J/Crl) purchased from Charles River were injected subcutaneously with 5 x 106 cells in 1:1 solution with Matrigel in one flank. Mice with established A549 tumors (100mm3 or 300mm3) were treated by oral gavage once daily with (50mg/Kg or 100mg/Kg) SD-066-4 dissolved at 20mg/ml in PEG300 and mixed 1:1 with 20% captisol or vehicle control. Tumors were measured daily by caliper. When tumor volume reached 1000mm3 animals were euthanized and tumors were harvested.

Immunofluorescence of Xenograft tumors

Tumors were embedded in O.C.T compound (Tissue-Tek) and frozen on dry ice. Frozen blocks were sectioned at -20 C˚. Slides with 10µm sections were fixed in formalin for 30 minutes then blocked in 5% BSA in TBST. Primary antibodies were diluted 1:200 in 5% BSA in TBST and incubated overnight at 4C˚. Slides were washed in TBST and incubated with secondary antibody diluted in 5% BSA for one hour at room temperature. Slides were washed in TBST and mounted in Fluoromount-G.

S-Acyl Exchange-PLA

Slides with 10µm sections were fixed in formalin for 30 minutes then dehydrated in xylene and rehydrated in a series of ethanol washes. Antigen retrieval was performed using citrate buffer (Electron Microscopy Sciences #62706-13). Immediately after antigen retrieval, the slide was washed with ABE buffer (150mM NaCl, 50mM Hepes pH 7.4, and 10mM EDTA, and 0.2%Triton X-100) and then incubated in NEM buffer (50mM NEM, 150mM NaCl, 50mM Hepes pH 7.4, and 10mM EDTA, and 0.2%Triton X-100) at 37oC for 1hour. Next, samples were washed one time in ABE buffer, followed by incubation in NEM buffer overnight at 4C˚. The samples were washed three times in ABE buffer followed by one hour incubation with ABE buffer containing hydroxylamine (0.7M) as described above. The slides were then washed once in ABE buffer followed by three washes in DPBS. The samples were incubated with DPBS containing 10µM iodoacetamide-alkyne, followed by washing in DPBS. Samples were treated with a click reaction mix containing 25µM OG 488 Azide (Click Chemistry Tools #1264-1), 1mM $CuSO_4$, and 1mM tris(2-carboxyethyl) phosphine (TCEP) in DPBS for 45-60 minutes at room temperature in the dark then blocked in 5% BSA in TBST. Incubation with primary antibodies specific for phosphorylated EGFR (Y1068) (Novus Biologicals #MAB3570) and for fluorescein/OG 488 (Life Technologies A-889) was done overnight at 4oC. The Duolink assay was performed as per manufacturer's protocol. Samples were imaged on 40x magnification and Z-series were captured and deconvolved using Leica software. Deconvolved images were quantified for the number of PLA puncta per field.

Cell Viability

Cells were treated with DMSO or SD-066-4 at specified concentration for 72hrs and viable cell number was measured by Trypan Blue staining. Statistical significance was determined using 1-way ANOVA with Tukey's multiple comparison post-test.

Statistics

Statistical analyses were performed using Prism software version 7.0 (GraphPad). Experiments are reported as mean ± s.e.m as noted in the legends. Data were analyzed using a 2-tailed Student's t-test for comparison between 2 data sets. Multiple comparisons were analyzed by 2-way ANOVA, followed by Tukey's multiple-comparison correction. A P-value of less than 0.05 was considered statistically significant. Immunoblot experiments were performed 3 times. Drug response curves and cell viability experiments were performed 3 times with 3 replicates each for each dose.

Study Approval

All experiments involving live animals were performed in compliance with the guidelines set forth in the Public Health Service Policy on the Humane Care and Use of Laboratory Animals. Mice were housed in a pathogen-free facility at the American Association for Laboratory Animal Science–accredited Animal Facility at the University of Pennsylvania Perelman School of Medicine. All studies were performed under protocols approved by the Institutional Animal Care and Use Committee (IACUC) at the University of Pennsylvania Perelman School of Medicine (#804774).

**Supplementary Figures**

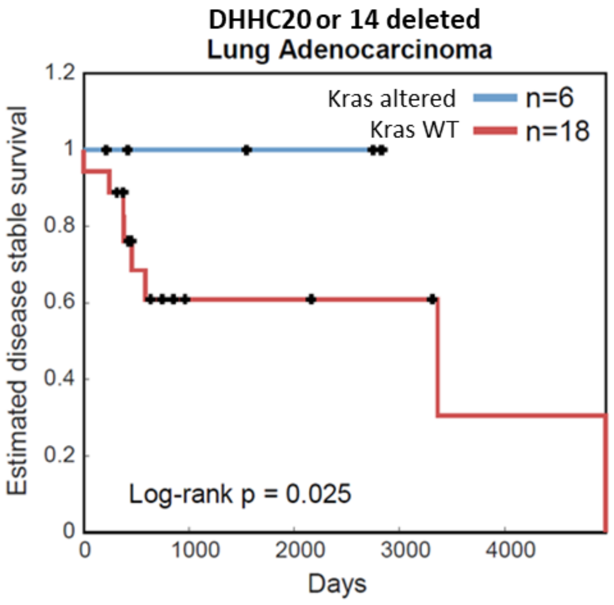


**Fig. S1.**

Lung adenocarcinoma patients with deletions in ZDHHC20 or ZDHHC14. When paired with Kras alterations (blue) patients have a significant increase in Estimated disease stable survival compared to patients with wild type Kras (red) (TCGA database) Log-rank Test 0.025.

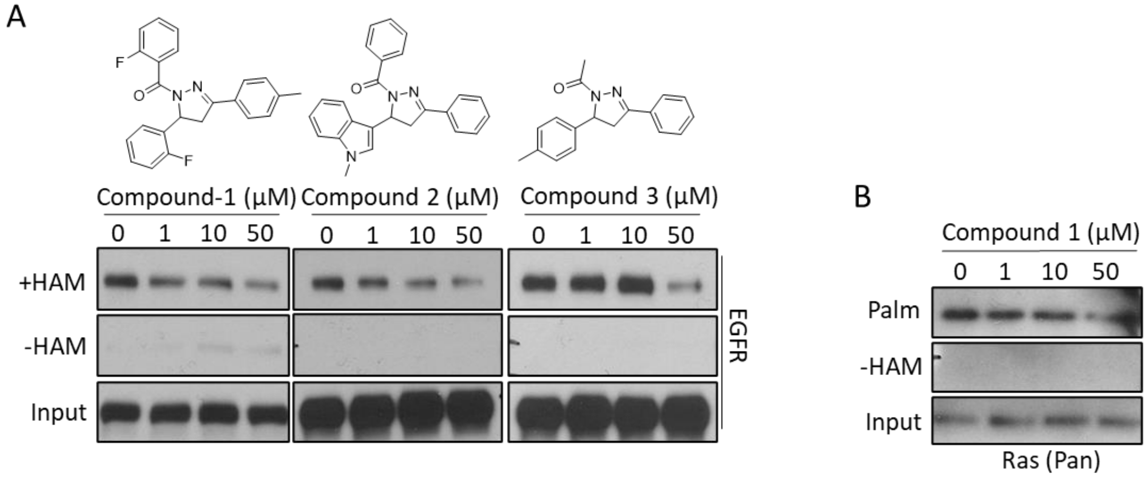


**Fig. S2.**

(**A**) ROCS identified compounds 1-3 inhibit EGFR S-acylation after one hour of treatment with increasing dosage. (**B**) Ras S-acylation after treatment with compound 1.

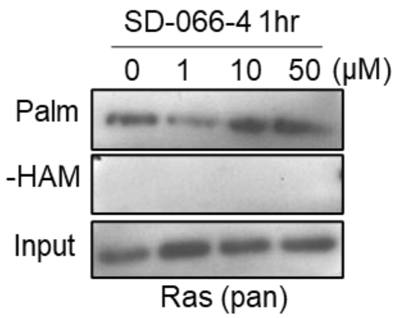


**Fig. S3.**
SD-066-4 is unable to inhibit Ras S-acylation.

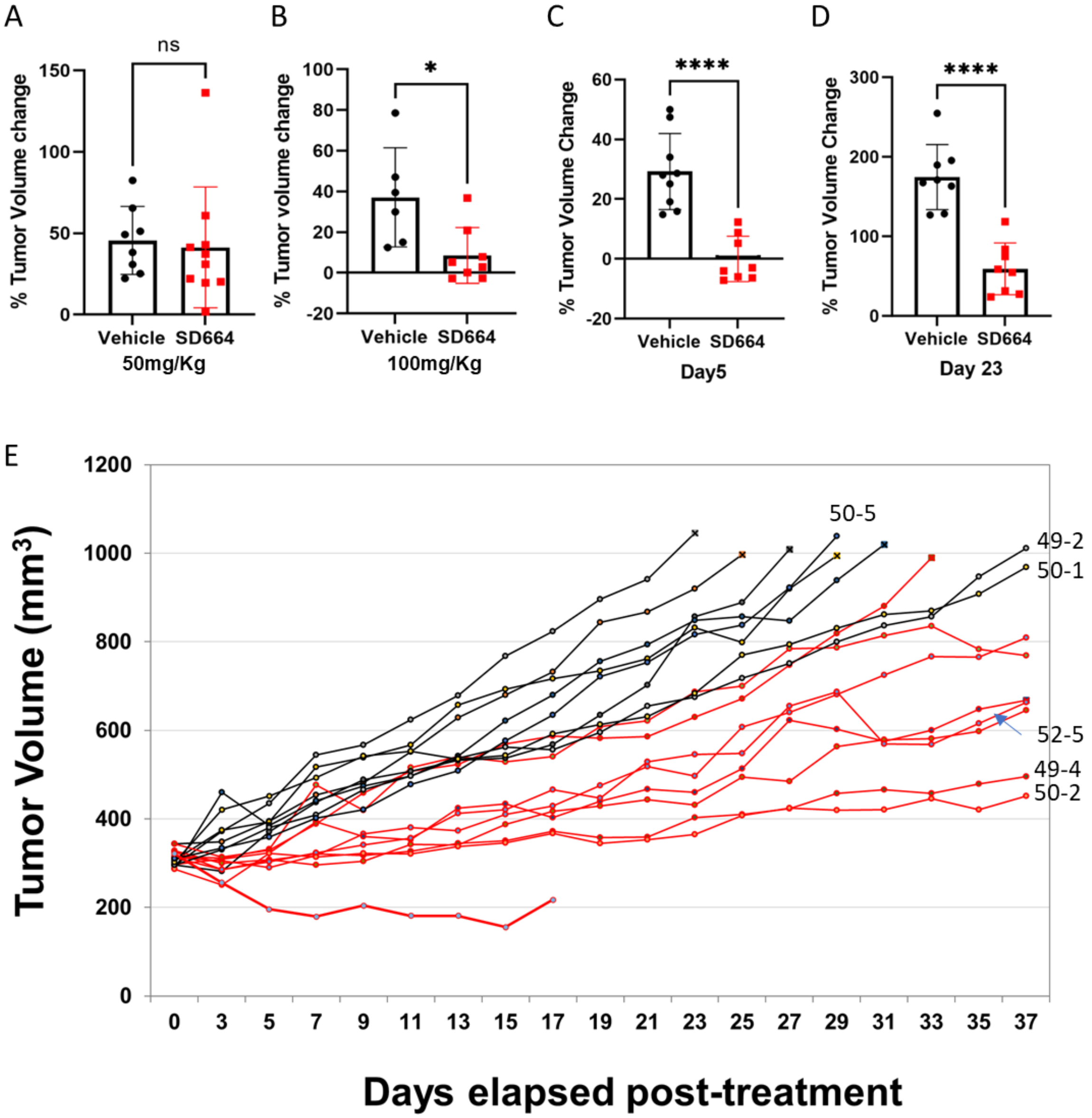


**Fig. S4**

(**A**) Treatment of xenograft A549 tumors with 50mg/kg SD-066-4 had no effect on tumor growth. (**B**) Increasing SD-066-4 dosage to 100mg/kg inhibited tumor growth. (**C**, **D**) Change in volume of A549 LUAD xenograft tumors on Day 5 and Day 23 of vehicle vs. SD-066-4 treated tumors. Treatment was initiated when tumors were 300mm3. Student's T-test. (**E**) Xenograft tumors treated with 100mg/kg or vehicle control were measured daily over 37 days. Individual tumors that were processed for immunofluorescence staining and SAE-PLA assays are labeled. One tumor regressed and the animal was euthanized on day 17 to conserve compound. The tumor was excluded from the average percent change in tumor volume in Figure 5.

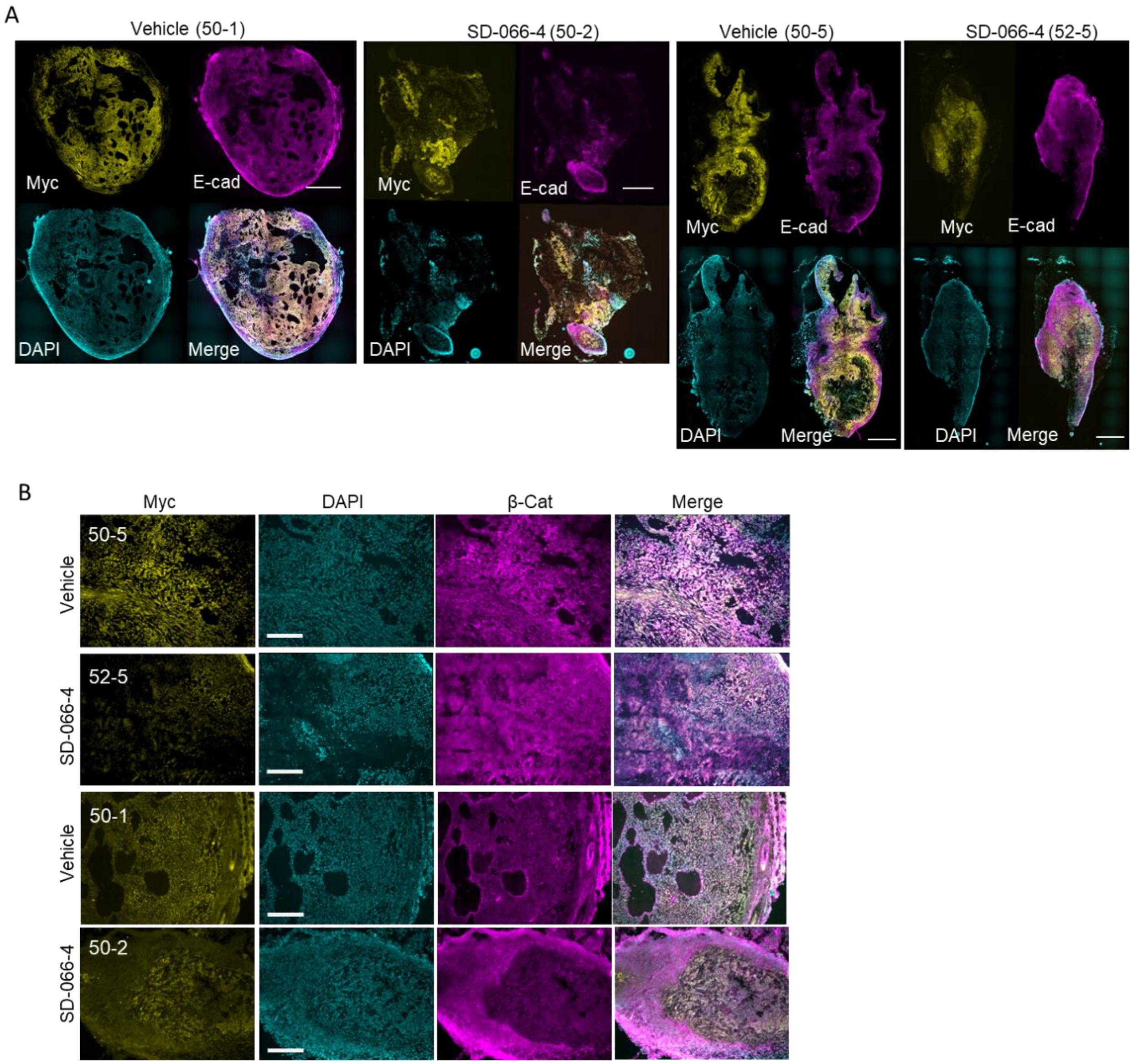


**Fig. S5**
(**A**) Immunofluorescence staining of SD-066-4 treated, and vehicle treated xenograft LUAD tumors. Anti-Myc (yellow) anti-E-cadherin (magenta), DAPI (cyan). Tile scan at 10X magnification (Scale bar = 1mm). (**B**) Immunofluorescence staining of frozen tumor sections probed with anti-Myc (yellow) anti-β-catenin (magenta) and DAPI (cyan). 40X magnification, (Scale bar = 50µm).

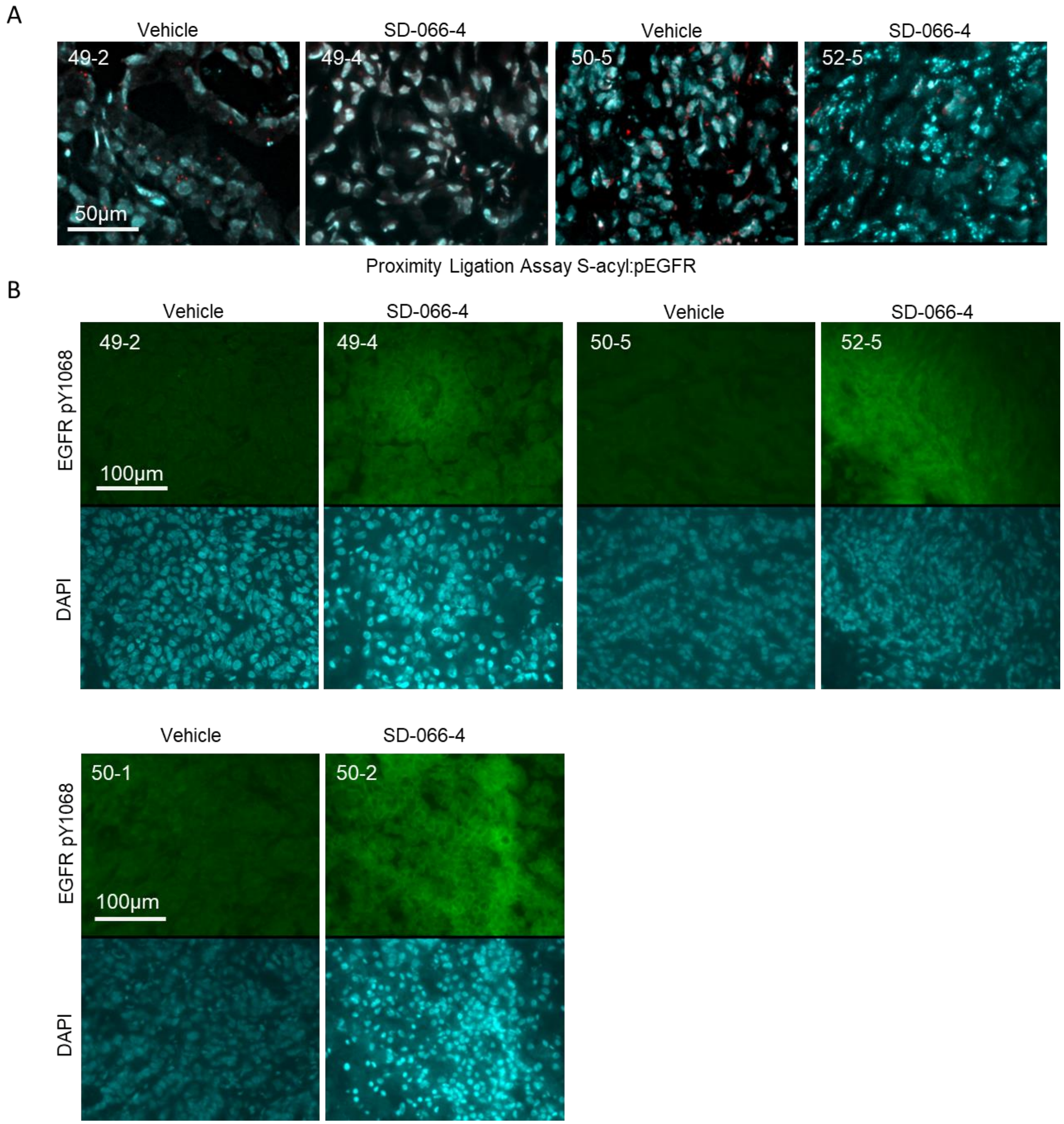


**Fig. S6**
(**A**) S-acyl exchange-PLA assay detects dually phosphorylated and S-acylated EGFR. S-acylated pEGFR is reduced in SD-066-4 treated tumors (Scale bar = 50μm). (**B**) pEGFR levels are increased with SD-066-4 treatment (Scale bar = 100μm).

| Gene | # Unique Peptides | T/C Ratio | P Value |
|---|---|---|---|
| SEC23A | 4 | 0.350522 | 0.046757 |
| DDOST | 5 | 0.392186 | 0.013666 |
| CD82 | 4 | 0.39437 | 0.00889 |
| PDIA5 | 4 | 0.405375 | 0.003615 |
| SORD | 4 | 0.421462 | 0.010768 |
| SF3A3 | 4 | 0.428277 | 0.017667 |
| CAPRIN1 | 4 | 0.472313 | 0.023842 |
| AP2B1 | 8 | 0.495759 | 0.009243 |
| CEMIP2 | 4 | 0.500842 | 0.014445 |
| SEPTIN2 | 3 | 0.534778 | 0.030022 |
| NUP93 | 7 | 0.550896 | 0.035657 |
| DDX19B | 6 | 0.558608 | 0.006687 |
| RBM28 | 7 | 0.569476 | 0.031771 |
| ARL13B | 5 | 0.570741 | 0.01786 |
| SSB | 7 | 0.572293 | 0.0181 |
| POP1 | 9 | 0.583062 | 0.019886 |
| **EGFR** | **10** | **0.59251** | **0.035717** |
| SEC24C | 7 | 0.597069 | 0.004796 |
| RPS21 | 3 | 0.608512 | 0.034804 |
| HECTD1 | 23 | 0.614295 | 0.018073 |
| COX5B | 3 | 0.618329 | 0.000983 |
| GNB1 | 5 | 0.620675 | 0.039643 |
| NAXE | 3 | 0.621208 | 0.046812 |
| NUP160 | 6 | 0.633837 | 0.034299 |
| NUP205 | 14 | 0.634219 | 0.034502 |
| USP7 | 10 | 0.648248 | 0.007483 |
| ACTBL2 | 4 | 0.65578 | 0.006051 |
| RPL9 | 5 | 0.665029 | 0.022296 |
| GOLIM4 | 13 | 0.666568 | 0.036589 |
| GTPBP4 | 15 | 0.667526 | 0.044498 |

**Table S1.**

Proteins from DMSO and SD-066-4 treated NCI-H23 cells were enriched by ABE and analyzed by LC-MS/MS. Ratio of Treated/Untreated (SD-066-4/DMSO).
